# Maternal n-3 enriched diet reprograms neurovascular transcriptome and blunts inflammation in neonate

**DOI:** 10.1101/2024.01.22.576634

**Authors:** Tetyana Chumak, Amandine Jullienne, C. Joakim Ek, Maryam Ardalan, Pernilla Svedin, Ryan Quan, Arjang Salehi, Sirus Salari, Andre Obenaus, Zinaida S Vexler, Carina Mallard

## Abstract

Infection during perinatal period can adversely affect brain development, predispose infants to ischemic stroke and have lifelong consequences. We previously demonstrated that diet enriched in n-3 polyunsaturated fatty acids (PUFA) transforms brain lipid composition and protects from neonatal stroke. Vasculature is a critical interface between blood and brain providing a barrier to systemic infection. Here we examined whether maternal PUFA-enriched diets exert reprograming of endothelial cell signalling in 9-day old mice after endotoxin (LPS)-induced infection. Transcriptome analysis was performed on brain microvessels from pups born to dams maintained on 3 diets: standard, n-3 or n-6 enriched. N-3 diet enabled higher immune reactivity in brain vasculature, while preventing imbalance of cell cycle regulation and extracellular matrix cascades that accompanied inflammatory response in standard diet. LPS response in blood and brain was blunted in n-3 offspring. Cerebral angioarchitecture analysis revealed modified vessel complexity after LPS. Thus, n-3-enriched maternal diet partially prevents imbalance in homeostatic processes and alters inflammation rather than affects brain vascularization during early life. Importantly, maternal diet may presage offspring neurovascular outcomes later in life.

## Introduction

Perinatal infection can impair neurodevelopment being associated with motor and cognitive abnormalities in infants (Sewell, Roberts, & Mukhopadhyay, 2021) and predisposing infants to stroke. Pathogenesis and timeline differ in neonatal and adult stroke due, among other things, to differences in blood brain barrier (BBB) response (Fernández-López et al., 2012; Mallard, Ek, & Vexler, 2018). The BBB makes up a unique semipermeable interface between the systemic circulation and brain parenchyma. Endothelial cells, a major component of the BBB, ensure the delivery of oxygen and nutrients and enable crosstalk between blood and brain (Kadry, Noorani, & Cucullo, 2020). The brain vasculature continues to develop and refine itself in the postnatal period in synergy with the overall neurodevelopment and neuroplasticity (Coelho-Santos & Shih, 2020). Proper function of brain endothelial cells is crucial for brain development, as their genetic abnormality can cause long-term neurovascular and behavioural alterations (Ouellette et al., 2020). Endothelial cells function also as conditional immune cells expressing pattern recognition and cytokine receptors, presenting antigens and releasing cytokines in response to immune stimulation (Amersfoort, Eelen, & Carmeliet, 2022).

Long-chain polyunsaturated fatty acids (PUFAs) are critical components of brain cell membranes. Both n-3 and n-6 long-chain PUFAs are highly enriched in the brain and are mainly represented by docosahexaenoic acid (DHA, 22:6 n-3) and arachidonic acid (AA, 20:4 n-6) respectively. High n-3 PUFA maternal status has beneficial effect on pregnancy duration, fetal brain development and eases neurovascular complications of premature birth (Carlson et al., 2013; Gustafson et al., 2022; Hellstrom, Nilsson, et al., 2021; Morton et al., 2020; Tam et al., 2016). PUFA content in the cell membranes not only modulates mechanistic membrane properties but also defines the nature of surrounding environment: bioactive lipid derivatives of PUFAs are thought to contribute to creation of a pro-(n-6) or anti-(n-3) inflammatory milieu. Modulatory effects of n-3 PUFAs on immune function of various immune cells such as macrophages or microglia have been studied (Gutierrez, Svahn, & Johansson, 2019). We previously demonstrated that diet enriched in n-3 PUFA transforms brain lipid composition in the offspring and protects the neonatal brain from stroke, in part by blunting injurious immune responses (Chumak et al., 2022). However, how PUFAs affect the functions of brain endothelial cells in response to an inflammatory stimulus in the neonatal period is unknown.

In the current study, we aimed to characterize the cerebral endothelial transcriptome under physiological conditions (unperturbed mice) and following endotoxin (LPS) administration. Furthermore, we evaluated impact of different maternal PUFA diets on LPS-induced changes of endothelial RNA transcripts, systemic and cerebral inflammation, and angioarchitecture in the offspring.

## Methods

### Animals

Three- to 6-week-old C57BL/6J mice were purchased from Charles River Laboratories (Sulzfeld, Germany) and housed at the Laboratory for Experimental Biomedicine (University of Gothenburg, Gothenburg, Sweden), or purchased from Jackson Laboratories (USA) and housed at the Gillespie Neuroscience Research Facility vivarium (University of California, Irvine). Mice were provided with nesting material, shelters and ad libitum access to food and water. All experiments were performed in accordance with ethical protocols approved by the Gothenburg Animal Ethics Committee (no. 32/2016 and 663/2018) and by the University of California Irvine Institutional Animal Care and Use Committee (no. 18-127).

### Diet and treatment

Breeding cages were randomly assigned to one of the experimental isocaloric diets (AIN-93G, BioServ): standard (Stand, BioServ #S7628), n-3 long chain (LC)-PUFA enriched (n-3, BioServ #S7630; soybean oil replaced with Menhaden Fish oil) or n-6 LC-PUFA supplemented (n-6, BioServ #S7629; based on corn oil). The detailed composition of the diets is presented in our previous publication (Chumak et al., 2022). The day of birth was defined as postnatal day (P) 0. At P9 pups received a single intraperitoneal injection of lipopolysaccharide (LPS from E. coli O55:B5, 1mg/kg, Cat# 423, List labs) or equivalent volume of saline. The dams remained on the experimental diet until offspring sacrifice (P10-P12). Efforts were taken to include equal number of pups of both sexes in each experimental group.

### Microvessel isolation and RNA sequencing

Microvessels were isolated from mice brains at 24 hours following LPS/saline treatment. Brains from three pups were pooled together to obtain one sample. Animals were gently perfused with cold 0.9% saline, cerebellum and olfactory bulbs were removed, and the remaining brain hemispheres were transferred to a petri dish with cold Ringers-Hepes (RH) buffer kept on ice, where meninges were carefully removed. Brain tissue was then homogenized in RH solution containing 1% albumin using a 7ml Dounce homogeniser with one gentle stroke. All following steps were conducted in a cold room using cold solutions. The homogenate was centrifuged at 500 g for 10 min and supernatant removed. The pellet was thoroughly mixed with 7mL of 17.5% 70kDa dextran solution by alternative aspiration/expulsion and centrifuged at 3000g for 40 min. The supernatant was removed, and pellet resuspended in 1% albumin RH solution. The solution was then filtrated drop-by-drop through a 100µm mesh, and microvessels collected on a 20 µm mesh. The vessels were resuspended in RH containing 0.1% albumin and spun at 500 g to form a microvessel pellet which was frozen at -80. Total RNA was extracted using the miRNeasy Micro Kit as per the guidelines from the manufacturer. RNA amount was determined through qubit analysis (Invitrogen). Preparation of RNA library and transcriptome sequencing was carried out by Novogene Co., LTD (Cambridge, UK). Stranded RNA sequencing with sequencing depth 60M pair-end reads (2*250) was performed.

### Read alignment and gene expression quantification

FastQC (version 0.11.2) was used for quality control of FASTQ format files and trimming of the adapter content and quality trimming was performed using TrimGalore (version 0.4.0) tool, a wrapper around Cutadapt (version 1.9) (Martin, 2011). Mapping of trimmed reads were mapped to the Mus-musculus mm10 genome using STAR (version 2.5.2b) and quantification of reads mapping to each gene was performed using FeatureCounts (version 1.6.4) (Dobin et al., 2013; Liao, Smyth, & Shi, 2014). List of annotated genes was downloaded from Ensembl (version 83) Mus-musculus GRCm38 (Flicek et al., 2014).

### Differential gene expression analysis and bioinformatics

Differential expression analysis was performed using DESeq2 v1.32.0 from Bioconductor (Love, Huber, & Anders, 2014). For pairwise comparisons, we compared diets using standard diet as baseline and then LPS vs saline controls for each diet. Pathway analysis was done through the Qiagen Ingenuity Pathways Analysis (IPA) (QIAGEN Inc., https://digitalinsights.qiagen.com/IPA) using genes with differential expression for each diet. During pathway analysis and visualization of the results using IPA, closely related pathway terms were omitted.

REACTOME browser was used to functionally annotate the genes regulated by LPS across diets (Gillespie et al., 2022; Kramer, Green, Pollard, & Tugendreich, 2014). For this, lists of differentially expressed genes were analysed independently for each diet and the result of this overrepresentation analysis was graphically expressed in REACTOME maps. These maps highlight (in yellow) the biological processes regulated by LPS within each diet.

To explore the data, perform Principal component analysis (PCA) and generate heatmaps, we used Qlucore Omics Explorer (Qlucore AB, Lund, Sweden). For PCA plots the lists of genes annotated to immune system (1292 genes), cell cycle (593) and ECM organisation (280) were extracted from the REACTOME website and used as input.

Volcano plots were generated using GraphPad Prism version 10.0.3 (GraphPad Software, Boston, Massachusetts USA, www.graphpad.com).

### Cytokine Multiplex Assays

At P10 (24h after injection) or P12 (72h after injection) the pups were deeply anesthetized via intraperitoneal administration of pentobarbital (Cat#338327, APL), blood was collected through heart puncture, mixed with 5 μl of 100 mM ethylenediaminetetraacetic acid (EDTA, ED2P, Sigma-Aldrich) and centrifuged to obtain blood plasma for further analysis. Pups were then transcardially perfused with 0.9% saline and the brain hemispheres were collected. Flash frozen on dry ice plasma samples and brain hemispheres were stored at -80°C until further processing. Cytokine/chemokine levels were measured using Bio-Plex Pro Mouse Cytokine Standard 31-Plex kit (Cat# 12,009,159, BioRad) on a Bio-Plex 200 analyser as per the manufacturer’s guidelines. The baseline cytokine data has previously been published (Chumak et al., 2022).

To assess the cytokine response to LPS, for each diet we calculated reaction to LPS using the following formula: Reaction to LPS = cytokine concentration (LPS) - mean cytokine concentration (saline)

Obtained values were then compared between different diet groups.

### Vessel painting

Imaging and analysis of vessel painted brains was performed as previously described (Salehi et al., 2018). Mice were injected intraperitoneally with sodium nitroprusside (75 mg/kg) and heparin (20 mg/kg) 5 min before an intraperitoneal injection of ketamine/xylazine (90/10 mg/kg). Vessel painting was performed by injecting a solution of 1,1’-Dioctadecyl-3,3,3’,3’-Tetramethylindocarbocyanine Perchlorate or DiI (Cat# D282, Invitrogen, 0.3 mg/ml in phosphate buffer saline (PBS) containing 4% dextrose, total volume of 250 μl) into the left ventricle of the heart, followed by a 10 ml PBS flush and a 20 ml 4% paraformaldehyde (PFA) perfusion, using a peristaltic pump (5 ml/min). Brains were extracted, post-fixed in 4% PFA for 24h, washed, and stored at 4°C in PBS until imaging. Well-labelled brains were imaged axially using a fluorescence microscope (Keyence BZX810, Keyence Corp., Japan) with a 2x magnification. The middle cerebral artery (MCA) vessels were imaged using a 10x magnification. Images were analysed for axial brain area, vessel and junction density, and vessel length using the Angiotool software (Zudaire, Gambardella, Kurcz, & Vermeren, 2011). Vascular complexity was analysed using the ImageJ plugin “FracLac” (Karperien, Ahammer, & Jelinek, 2013).

### Statistical analyses

Statistical analysis and visualization of cytokine and vessel characteristic data were performed using GraphPad Prism version 10.0.3. Outliers in the cytokine analysis were removed using the ROUT method based on the False Discovery Rate and identification of the outliers from nonlinear regression. Outliers in the vessel painting analysis were removed using the interquartile range method. Normality of the data distribution was checked using Q-Q plot of the data. Homogeneity of variance was tested using Brown-Forsythe test. Two-way ANOVA was used to test the main effects of diet and LPS treatment on cytokine levels in blood and brain, and on vessel characteristics. For cytokine levels, Tukey’s multiple comparison test was used to assess differences between diets in the LPS treated groups only, since the analysis on saline treated pups has been previously reported (Chumak et al., 2022). Cytokine reaction to LPS was compared between the diet groups using one-way ANOVA with Fisher’s LSD post-hoc test for data with normal distribution and equal variances, Welch’s ANOVA with Unpaired t with Welch’s correction post hoc test when groups had unequal variances, and Kruskal-Wallis test followed by uncorrected Dunn’s post hoc test when data had not normal distribution. For vessel characteristics, Tukey’s multiple comparison test was used to assess differences between groups and Sidak’s multiple comparison test was used to assess differences between timepoints among each group. Chi^2^ analysis was used to assess potential differences in proportion of capillary bed staining of the M3 MCA area between groups. Differences were considered statistically significant at p < 0.05. To analyse pup weight data, unpaired t test was used to compare two groups or Welch’s ANOVA with Dunnett’s post-hoc test to compare three groups of animals. Data for normalized counts of brain cell markers and vessel characteristics are presented as median with quartiles and at 10–90th percentile, horizontal bar in graphs for cytokine reaction to LPS represents median value. Violin plots are used for the pup weight change and brain area data. Graphs also include individual data points. Principal component analysis (PCA) and heatmaps were generated using Qlucore Software (Lund, Sweden).

## Results

### LPS-induced deficits in body weight gain persist longer in pups from dams fed n-6 diet

LPS did not result in pup mortality in any of the diets but significantly decreased weight gain in pups across all diets at 24 h compared to saline controls (Suppl. Fig. 1A). At 72 h after LPS treatment only pups fed n-6 diet exhibited significant deficiency in weight gain (Suppl. Fig. 1B). Consistent with our previous observations (Chumak et al., 2022), diet did not affect the pregnancy progress or number of pups born per litter but weight of the pups from the dams fed n-3 diet was 9% higher than the weight of the pups from the dams on the other two diets at P9, the time of LPS injection (Stand.: 4.16 ± 0.53, n-3: 4.51 ± 0.57, n-6: 4.08 ± 0.4 g; p = 0.003, Welch’s ANOVA, Suppl. Fig. 1C). Axial brain area was derived from the Angiotool analysis and revealed no differences between saline and LPS-treated groups at 24 h (Suppl. Fig. 1D) or 72 h (Suppl. Fig 1E).

### Brain vessel isolation procedure leads to highly enriched population of endothelial cells and pericytes

We first assessed purity of the isolated vessels using the normalized counts (transcripts per million) of various brain cell markers for endothelial cells, pericytes, astrocytes, microglia, neurons and oligodendrocytes, as summarised in Figure 1A. Transcripts for brain endothelial-associated proteins such as *Cldn-5*, *Glut1* and *Abcb1/p-gp,* showed the highest number of counts among the cell markers. The second highest counts were found for the pericyte-associated transcripts, such as *Pdgfrb* and *Cspg4/Ng2,* but with an average of 5-fold lower counts than the endothelia-associated genes. Forty-fifty-fold lower counts were observed for astrocyte-, microglia-, oligodendrocyte- and neuron-associated transcripts. Thus, the utilized isolation procedure led to a highly enriched population of endothelial cells with associated pericytes.

**Figure 1.**
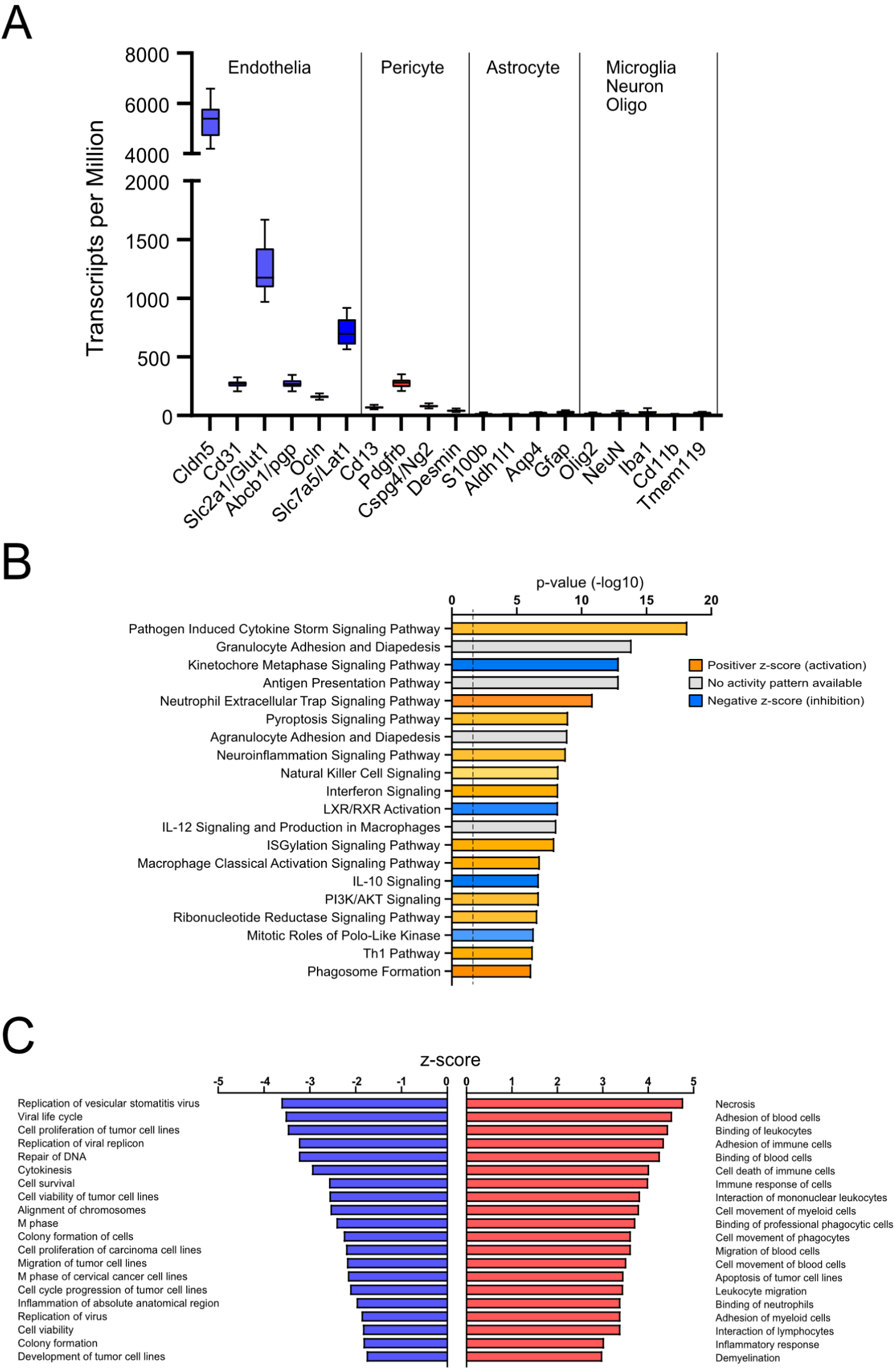
LPS leads to upregulation of inflammatory pathways while downregulating cell division and proliferation pathways in brain microvessels of offspring on standard diet. (A) Normalised counts of markers for endothelial cells (*Cldn-5, CD31, Glut1, Abcb1/p-gp, Ocln, Zo-1*), astrocytes (*Gfap, Aldh1h1, S100b, Aqp4*), pericytes (*CD13, Desmin, Pdgfrb, Cspg4/Ng2*), neurons (*NeuN*), oligodendrocytes (*Olig-2*), and microglia (*Iba1, CD11b*). (B) Top 20 canonical pathways regulated by LPS in standard diet group sorted based on overrepresentation p-value. The colours represent levels of the activation z-score: blue indicates downregulated and orange-upregulated pathway, pathways for which the direction of change is not possible to predict are shown in grey. (C) Top 20 up- and downregulated by LPS biological functions in brain microvessels in standard diet group sorted based on the z-score. Positive z-score indicates activation (red), negative z-score indicates inhibition (blue) of the biological function.

### LPS affects brain microvasculature transcriptome in a diet dependent manner

#### Standard diet

Analysis of gene transcripts in brain microvessels after saline or LPS injection in pups from the dams fed standard diet revealed 791 differentially expressed genes (DEGs). Utilizing IPA, we generated a list of top 20 canonical pathways sorted based on overrepresentation p-value using all DEGs (Fig. 1B). Most of the highest overrepresented pathways were related to innate immunity and inflammation with the majority showing significant positive activation score (z-score>+2) including: neutrophil extracellular trap (+5.38), macrophage classical activation (+3.71), neuroinflammation (+3.4), pathogen induced cytokine storm (+3.18), pyroptosis (+3.3) (Fig. 1B, Suppl. Table 1). No activity pattern was predicted for antigen presentation, granulocyte and agranulocyte adhesion and diapedesis signalling pathways (Fig. 1B, Suppl. Table 1).

To gain deeper insight and explore the biological processes that are activated or inhibited in brain microvessels after LPS injection in standard diet group, we also examined the biological functions generated by IPA. In the Fig. 1C the top 20 positive and negative biological processes are listed sorted by z-score. Among the top positive z-scores, many are related to cell death (necrosis: +4.8; cell death of immune cells: +4.0; apoptosis of tumour cell lines: +3.5) and inflammatory response, particularly associated with the interaction of circulating immune cells with the vessel wall (cell movement: +3.5; migration: +3,6; adhesion: +4.5 and binding: +4.3 of blood cells etc.; Fig. 1C, Suppl. Table 1). The top negative z-score functions were related to pathogen survival (viral life cycle: -3.5; replication of virus: -1.9, etc.), host cell survival (cell survival: -2.6 and viability: -1.8;), division (M phase: -2.42; alignment of chromosomes: -2.55; repair of DNA: -3.24; cytokinesis: -2.95; cell cycle progression: -2.12) and proliferation (cell proliferation: -3.5 and development: -1.76 of tumour cell lines; Fig. 1C, Suppl. Table 1). Thus, in brain microvessels of LPS-treated mice from dams fed a standard diet, upregulation of inflammatory response genes was accompanied by activation of cell death and downregulation of cell proliferation pathways.

#### PUFA diets

Comparisons of gene expression under physiological conditions showed no differences between diets, whereas LPS modulated gene expression in microvessels in diet-specific ways, as shown on a Venn diagram (Fig. 2A), a heatmap of normalized counts of all DEGs (Fig. 2B) and on volcano plots (Fig. 2C). In the Venn diagram and the heatmap, we combined all genes regulated by LPS (DEGs) within each diet (1214 genes in total). The total number of genes regulated by LPS was lower in PUFA diet groups than in mice on standard diet (742 in n-3 and 677 in n-6). Out of all DEGs, 379 were common between all three diets, and 325 were regulated by LPS exclusively in the standard diet group (Fig. 2A). Brain microvessels from n-3 and n-6 diets exhibited a higher number of overlapping DEGs (151) compared to the standard diet (53 in n-3 and 34 in n-6), which suggests a similar response to LPS. There was a lower proportion of downregulated genes in the n-3 (12% of DEGs) and n-6 (10%) diets compared to the standard diet (39%).

**Figure 2.**
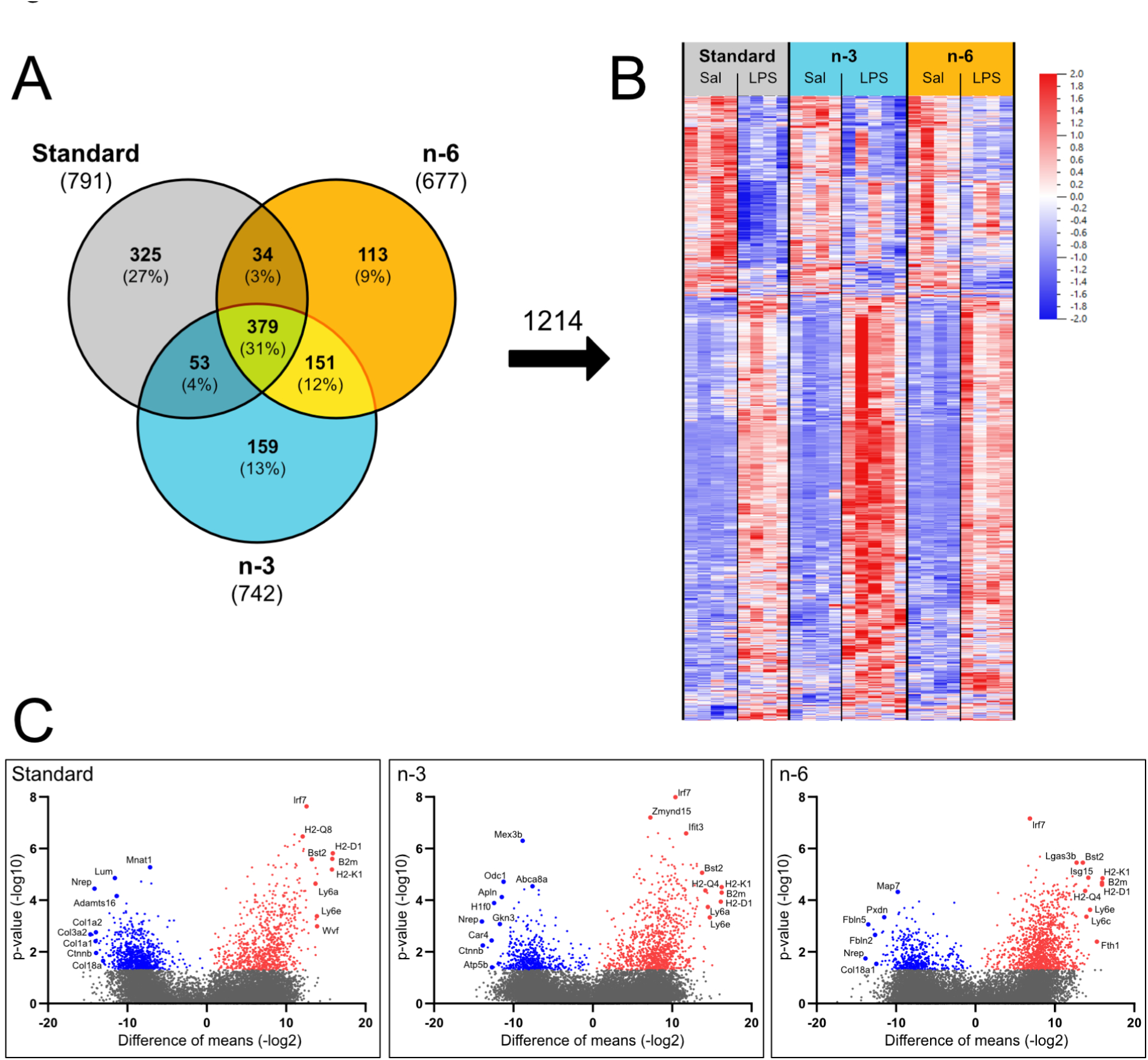
The most significantly upregulated genes in brain microvessels after LPS are related to inflammation in all diets, while ECM genes are among the most significantly downregulated genes in standard and n-6 diets only. (A) Venn diagram of all genes regulated by LPS (adjusted p < 0.05) within each diet. (B) Heatmap of normalized counts of all genes regulated by LPS across all diets (1214 genes in total) sorted using hierarchical clustering. Each column refers to one microvessel sample and each row shows level of a specific gene. Numbers of transcripts for each gene were normalized over all samples using z-score normalization and is shown using a colour scheme based on z-score distribution from -2 to 2. (C) Scatter plot (Volcano plot) of p-value (y-axis) versus difference of means (x-axis) of all genes within each diet. Significantly regulated genes are coloured blue (downregulated) or red (upregulated). Note the higher number of downregulated genes in the standard diet than in the n-3 or n-6 diets. Genes with high magnitude of change and low p-value have been highlighted.

Heatmap shows an overall more prominent upregulation of genes by LPS in the n-3 diet group, but more pronounced downregulation of some genes in the standard diet pups (Fig. 2B). Upregulated genes (labelled in volcano plot) are consistent across all diets with the majority associated with the immune system function (Fig. 2C). Several of the most downregulated microvessel mRNAs from pups on the standard diet were related to the extracellular matrix (ECM) (Lum, Adamts16) and collagen production (Col1a1, Col1a2, Col3a2, Col18a). The Col18a1 gene and some ECM genes (Pxdn, Fbln2, Fbln5) were also down-regulated in n-6 diet group. Among these, peroxidasin (Pxdn) and fabulin-5 (Fbln5) have previously been shown to be involved in angiogenesis (Chan et al., 2016; Medfai et al., 2019). The most downregulated transcripts in n-3 diet included several cell proliferation genes (H1f0, Gkn3, Odc1), but no ECM-related genes.

The REACTOME browser was used to functionally annotate and further compare the genes regulated by LPS across diets. In the REACTOME map for standard diet in Figure 3A, the three main categories identified above are highlighted (in yellow): immune system (innate-, adaptive-immunity and cytokine production), mitotic cell cycle (M phase etc.) and extracellular matrix organisation (ECM proteoglycans, collagen formation and degradation, and syndecan interactions). To a lesser extent, cell death and hemostasis categories were also identified. In contrast, following LPS, only immune system genes were strongly identified in n-3-fed animals, whereas n-6 diet exhibited a high overrepresentation of immune system, along with a minor representation of ECM genes, but no overrepresentation of cell cycle-related genes. The lists of pathways enriched by LPS across diets are presented in the Suppl. Table 2.

**Figure 3.**
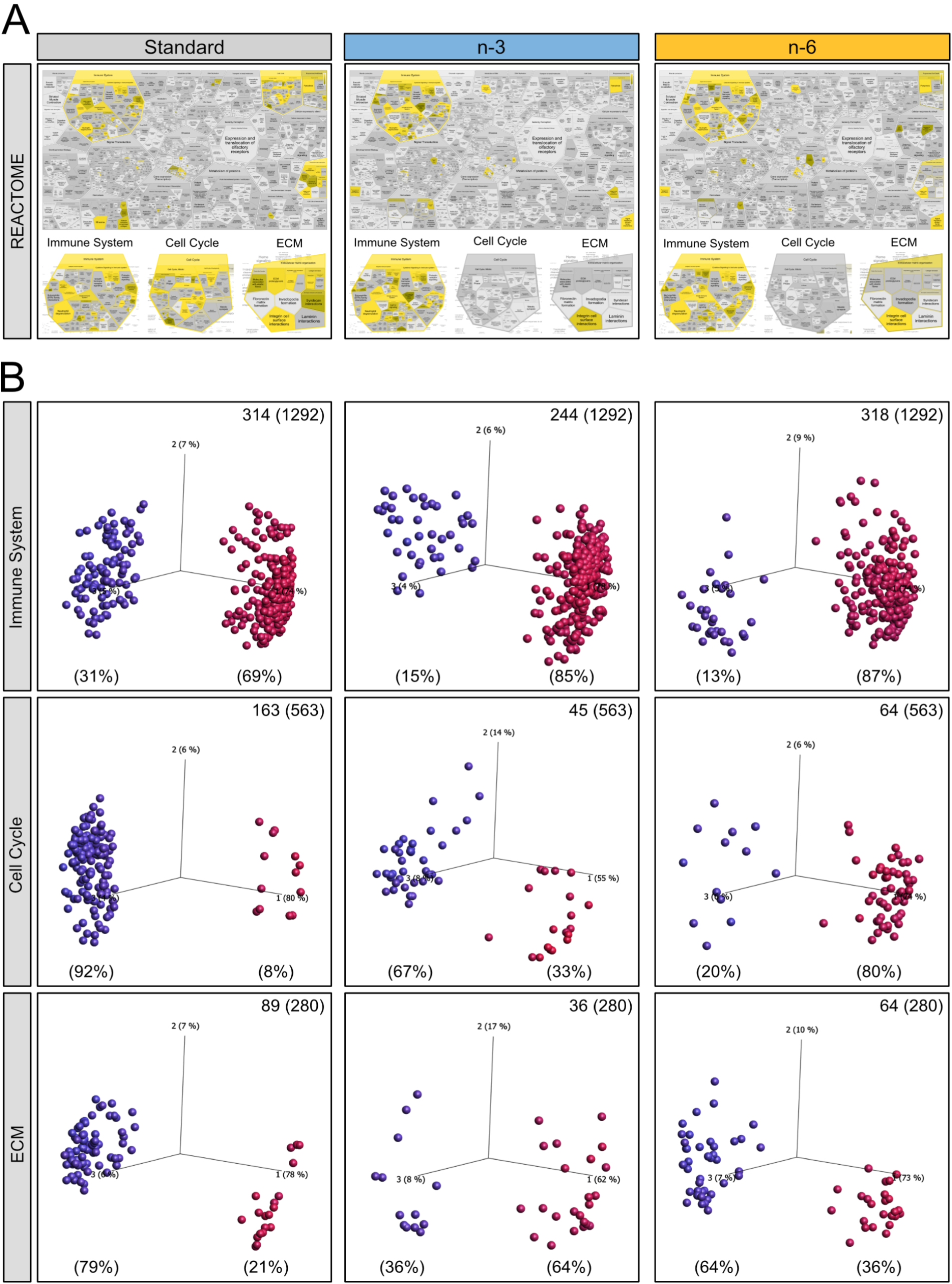
Functional annotation of genes validated stronger ECM gene response in standard and n-6 diet, additionally highlighting downregulation of cell cycle pathways only in standard diet. (A) Graphical pathway maps from overrepresentation analysis of all genes regulated by LPS within each diet created using the Reactome browser (adjusted p < 0.05; DeSeq). Yellow represents specific enrichment. Enrichment of pathways annotated to immune system (innate- and adaptive-immune system as well as cytokine response) is present in all three diets. Note high annotation to cell cycle and extracellular matrix organisation pathways in the standard diet. The n-3 and n-6 diet show no enrichment of cell cycle pathways and n-3 diet shows least enrichment of ECM pathways. (B) PCA plots of individual genes annotated to immune system (upper panel), cell cycle (middle panel), and extracellular matrix organisation (ECM, lower panel) regulated by LPS within each diet. Variables are colour coded according to distance between them rendering upregulated genes red and downregulated genes blue. Numbers of regulated genes are stated in upper right corner out of all analysed genes for each process (in brackets).

To explore directionality of LPS-induced gene changes within biological processes identified by REACTOME, we utilised variable PCA plots (Fig. 3B). Vast majority of genes associated with the immune system were upregulated in all diets; however, in n-3 and n-6 diets the percentage of downregulated immune genes was lower than in the standard diet group (Fig. 3B, upper panel). Complementing these findings, IPA analysis showed higher immune response (z-scores) in PUFA supplemented diets (Suppl. Table 1). PUFA diets show less regulation of cell cycle genes as well as less inhibition of these genes compared to the standard diet (Fig. 3, middle panel). IPA analysis concurred showing negative z-scores for cell division and proliferation only in standard diet while most of these biological functions were not overrepresented in PUFA diets (Suppl. Table 1). In addition, LPS activated cell death mechanisms in all diets while differently affecting expression of cell survival and viability genes: negatively in standard diet and positively in PUFA supplemented diets. PCA plots showed predominant inhibition of ECM genes in standard and n-6 diets, an effect not present in n-3 diet (Fig. 3, lower panel). Overall, the number of genes regulated in all these functions (immune system, cell cycle and ECM organisation) were highest in the standard diet and lowest in the n-3 diet group. Thus, PUFA supplemented diets diminished the effect of LPS on immune system and cell survival genes, while only n-3 diet reverted the LPS-triggered downregulation of ECM genes in standard diet.

Changes in cell survival, proliferation and impaired ECM function can result in BBB malfunction and altered angiogenesis (Krueger et al., 2015; Thomsen, Routhe, & Moos, 2017). However, we found very few cell junction genes that were regulated by LPS across all diets suggesting preserved BBB function (Suppl. Fig 2). In contrast, fewer angiogenic genes were downregulated in n-3 diet compared to n-6 and standard diets. IPA analysis showed activation of angiogenesis in n-3 diet (+2.25) while z-scores for n-6 (+1.70) and standard diet (+0.45) did not reach significance (z-score>2; Suppl. Table 1).

### N-3 diet diminishes and n-6 diet aggravates offspring plasma cytokine response to LPS

Differences in inflammatory signalling in brain microvessels in standard diet compared to PUFA supplemented diet groups prompted us examine the extent of systemic inflammatory changes on protein level in pups from dams fed different diets. See Figure 4A, Supplemental Figure 3A and Supplemental Tables 3, 4 for cytokine levels in blood of pups from different groups and the results of statistical analysis.

**Figure 4.**
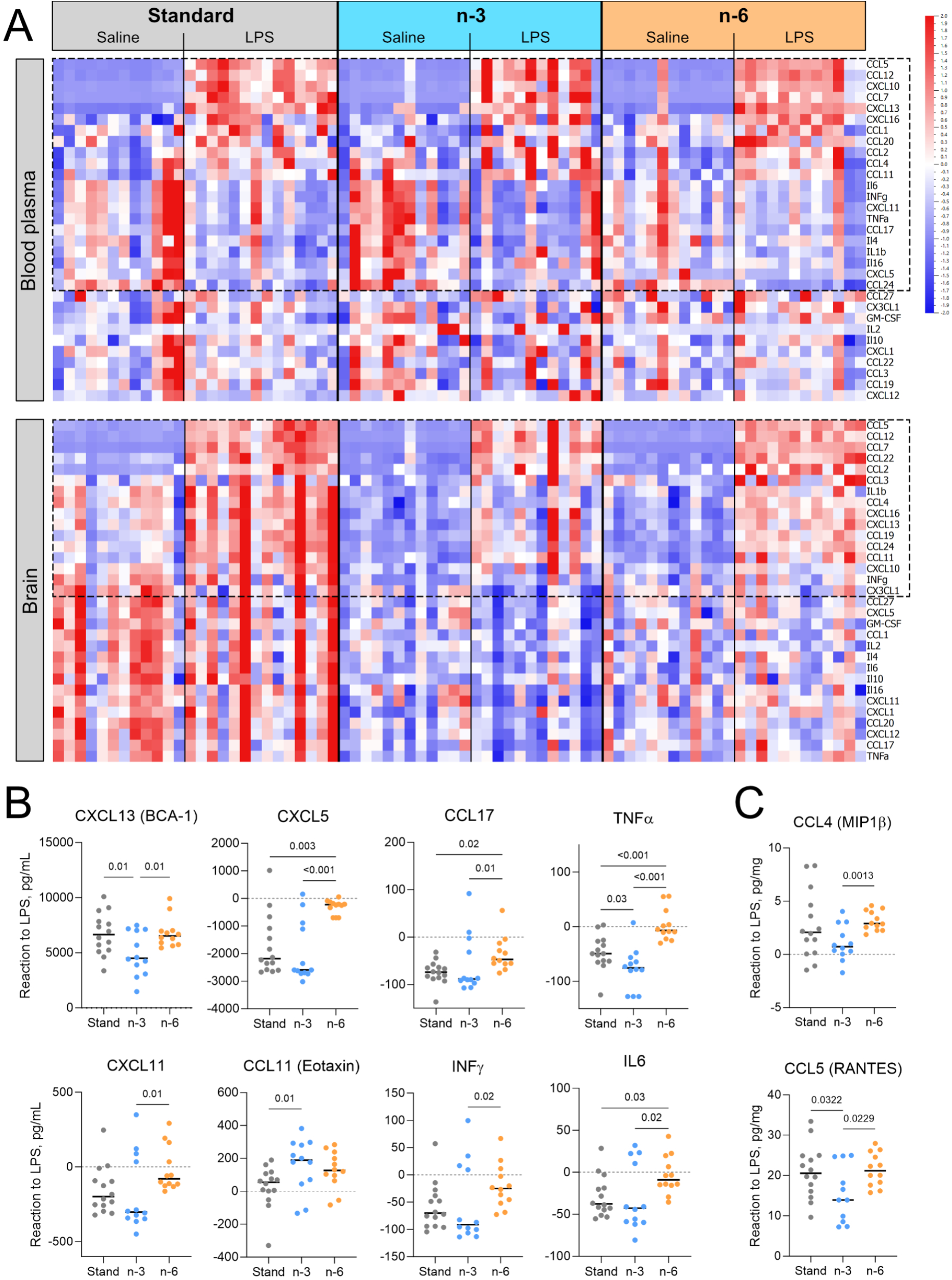
PUFA diets modulate cytokine levels in offspring blood and brain 24 h after LPS injection. Cytokine levels in the offspring blood plasma and brain 24h after LPS injection. (A) Heatmap of cytokine levels analysed in blood plasma and brain of offspring from dams fed the standard, the n-3 or the n-6 enriched diet and injected with saline or LPS at P9. Each column refers to one animal and each row shows levels of a specific cytokine. Each cytokine level was normalized over all samples using Z-score normalization and is shown using a colour scheme based on Z-score distribution from -2 to 2. Cytokines regulated by LPS (based on two-way ANOVA results) are highlighted with the frame. (B, C) Graphs showing the reaction to LPS in plasma (B) or brain (C). Zero level indicates no difference between the level of the cytokine in LPS sample and the mean level of the cytokine among the saline samples. Only graphs with significant differences between diet groups are presented. P values presented on the graphs show results of the post-hoc test. One-way ANOVA with Fisher’s LSD (B: CXCL13, TNFα; C: CCL5), Welch’s ANOVA with Unpaired t with Welch’s correction (C: CCL4) and Kruskal-Wallis with uncorrected Dunn’s (B: CXCL5, CCL17, CXCL11, CCL11, INFγ, IL6) post hoc tests.

Two-way ANOVA analysis showed a significant main effect of diet on the plasma levels of 4 cytokines at 24 h and 2 cytokines at 72 h after LPS injection (Suppl. Table 3, 4). Among them, Tukey’s post-hoc test showed higher post-LPS plasma level of CXCL12 24 h after LPS treatment in pups from dams fed n-6 diets compared to the standard diet group (Suppl. Table 3).

At 24 h after injection, significant main effect of LPS treatment on the level of 21 inflammatory cytokines/chemokines was found (Suppl. Table 3). Levels of 11 cytokines were higher and 10 cytokines were lower in plasma of LPS-treated animals compared to saline controls suggesting the ongoing process of resolution of systemic inflammation at this time point (Fig. 4A, upper panel; Suppl. Table 3). For these 21 cytokines we calculated the reaction to LPS and found differences (one-way ANOVA or Kruskal-Wallis) in cytokine response to LPS between the diets for CXCL13 (p = 0.019), CXCL5 (p < 0.001), CCL11 (p = 0.034), CXCL11 (p = 0.043), CCL17 (p = 0.02), TNFα (p < 0.001), INFγ (p = 0.048) and IL6 (p = 0.035), see Figure 4B. The reaction to LPS in the n-3 diet group was often downregulated compared to other diets consistent with a stronger anti-inflammatory response (Fig. 4B). In contrast, in the n-6 group, the response to LPS, was mostly less downregulated compared to standard and/or n-3 diet (Fig. 4B).

At 72 h after LPS injection, significant main effect of treatment on cytokine levels in plasma was found for 9 cytokines with the majority (8) being lower in LPS-exposed group compared to saline controls (Suppl. Fig. 3A, upper panel, Suppl. Table 4). Reaction to LPS was different between the diets (one-way ANOVA or Kruskal-Wallis test) for CCL24 (p = 0.038), IL4 (p = 0.0007), CCL7 (p = 0.034), CCL20 (p = 0.035), CCL17 (p < 0.0001), TNFα (p = 0.009), CCL22 (p < 0.0001), and CCL19 (p= 0.002). For most of these cytokines, the levels in n-6 diet group were the farthest from the baseline levels compared to standard and/or n-3 diets.

Taken together, analysis of systemic cytokine response suggests inflammatory response in blood plasma 24 h after LPS injection in all diets, with strongest anti-inflammatory effect 24 h and fastest restoration of the cytokine balance by 72 h after initiation of inflammation in n-3 diet. The opposite dynamics was found in n-6 diet.

### PUFA supplemented diets decrease post-LPS cytokine levels in the offspring brain

To examine neuroinflammation after systemic LPS injection, we analysed the levels of inflammatory cytokines in the brain tissue at 24 and 72 h (Fig. 4A, lower panel). As we have previously reported, most brain cytokines and chemokines levels were lower in n-3 and n-6 diet pups at P10 (24 h) compared to standard diet (Chumak et al., 2022).

A significant main effect of diet was found on the level of 26 cytokines at 24 h and 16 cytokines at 72 h after LPS injection (Suppl. Tables 5 and 6). Post-hoc test showed that all 26 cytokines regulated at 24 h, except CXCL10, were lower in n-3 diet pups than in standard diet group. Among them, levels of 10 cytokines were lower also in n-6 diet compared to standard diet. At 72 h after LPS injection, the level of 7 cytokines was lower in the brains of LPS-treated pups from n-6 diet fed dams compared to the standard diet group.

A significant main effect of LPS treatment was observed for the levels of 16 cytokines in the brain 24 h after injection (Suppl. Table 5). For all of them except CX3CL1, the levels in the LPS-treated group were higher than those in the control group suggesting ongoing inflammation in contrast to plasma (Fig. 4A, lower panel; Suppl. Table 6). Results from one-way ANOVA showed differences in reaction to LPS between diets for CCL4 (p= 0.005) and CCL5 (p = 0.043), with the lowest reaction to LPS in n-3 diet (Fig. 4C).

Two-way ANOVA analysis showed that LPS treatment resulted in regulation of 7 brain cytokines at 72 h (Suppl. Fig. 3A, lower panel). Levels of 4 cytokines were elevated and levels of 3 were decreased after LPS (Suppl. Table 6, Suppl. Fig. 3A, lower panel) in all diet groups suggesting resolution of inflammation in progress. Among them, the CCL22 reaction to LPS n-3 and n-6 diet (p < 0.0001, one-way ANOVA) and the CCL11 reaction to LPS in the n-3 diet (p = 0.0289, Kruskal-Wallis test) were lower compared to standard diet, see Suppl. Figure 3C.

A strong cytokine response was triggered in the neonatal brain at 24 h after LPS injection but it declined by 72 h. For some cytokines, this response was suppressed in the brains of pups from n-3 diet group.

### Brain vessels network is unaffected by diet or by LPS

Considering the notable downregulation of genes associated with the cell cycle and ECM in the standard diet, coupled with the upregulated angiogenesis in the n-3 diet, we tested if diets affect brain angioarchitecture following LPS exposure. We first assessed axial brain vessel effects of diet on saline-treated animals at both 24 and 72 h post-saline injection (Fig. 5A) by injecting fluorescent dye (DiI) and examining DiI-labelled vessels using the Angiotool software. We found no significant differences in vessel density, junction density, average and total vessel length (data not shown). No significant differences in these vessel characteristics from 24 and 72 h in the saline-treated animals were found (Fig. 5B-E). LPS exposure did not result in any significant changes in vessel features at 24 h (Fig. 5F-I). However, 72 h post-treatment, mice in the standard diet group treated with LPS trended to have an increased vessel density and average vessel length compared to saline treated mice (p = 0.0837 and p = 0.0684, respectively, two-way ANOVA Fig. 5J, L), but not in n-3 and n-6 diets.

**Figure 5.**
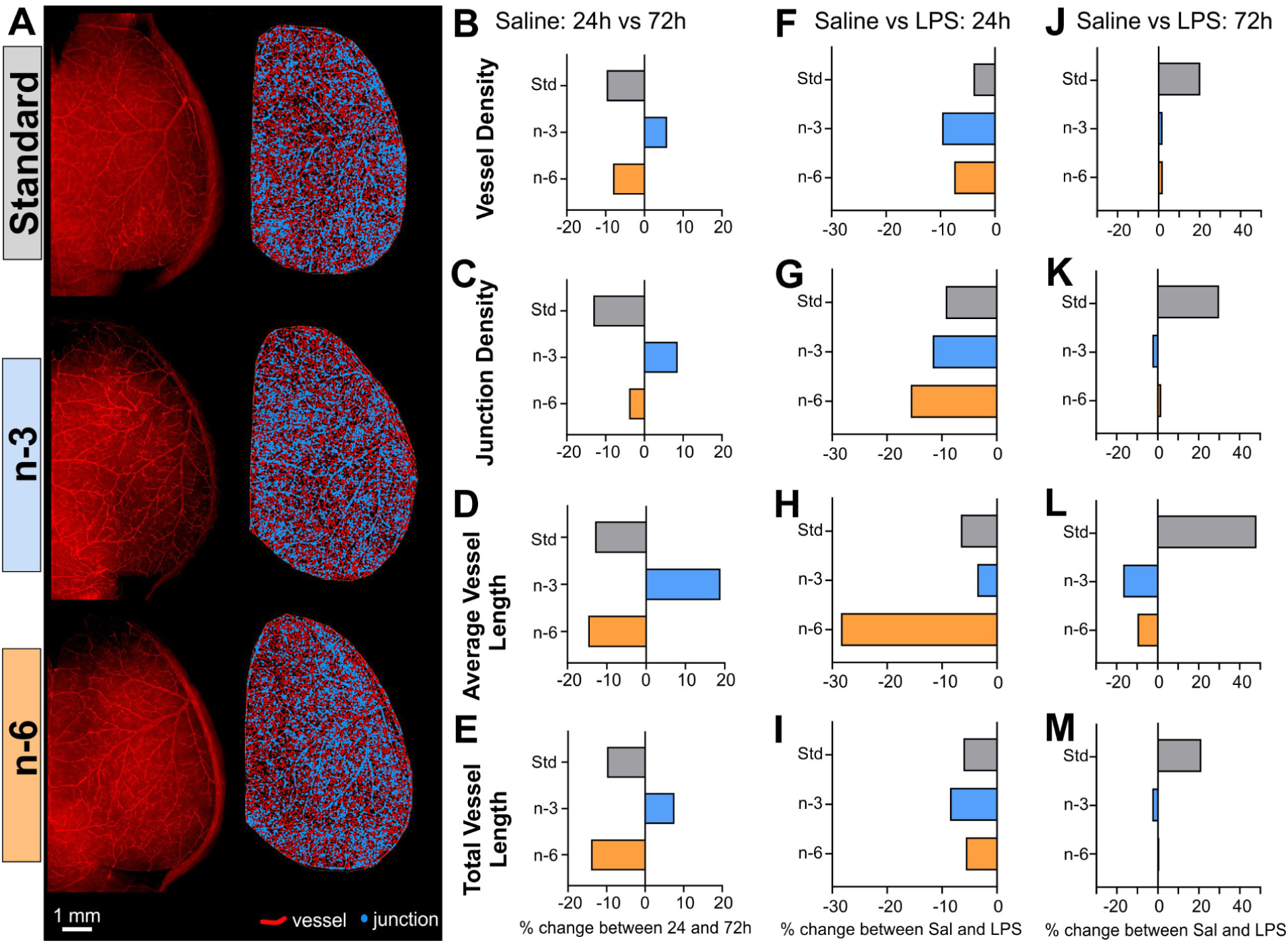
Morphological vessel characteristics are not affected by diet or treatment. (A) Left: Vessel painting image of one hemisphere from mice of the three diet groups treated with saline injection at 72 h. Right: Angiotool result images showing vessels in red and junctions in blue. Vessel density (B), junction density (C), average (D) and total vessel length (E) are not significantly affected by the diet nor by time in the saline condition. Similarly, vessel characteristics are not affected by LPS treatment at 24 h (F-I) nor 72 h (J-M).

Evaluation of the vessel network complexity using fractal analyses showed a global decrease in vessel local fractal dimension (LFD) (Fig. 6) consistent with decreased complexity (leftward shift, decreased skewness values) at 72 h (dotted lines) compared to 24 h (solid lines) in all groups. Skewness showed a significant time effect in the standard diet (p = 0.023, two-way ANOVA, Fig. 6B), n-6 diet (p = 0.012, two-way ANOVA, Fig. 6H) and a trending time effect for the n-3 diet (p = 0.058, two-way ANOVA, Fig. 6H). Skewness was significantly decreased in the LPS group in the standard and n-6 diets between 24 h and 72 h (Sidak’s test: p = 0.012 and p = 0.037, Fig. 6B and H) and trending in the n-3 diet (Sidak’s test: p = 0.098, Fig. 6E). Kurtosis of the LFD curves represents the number of vessels and a significant effect of time in the n-6 group (p = 0.030, two-way ANOVA, Fig. 6I) and again a trending effect of time in the n-3 group (p = 0.081, two-way ANOVA, Fig. 6F). No effect of time was found in the standard diet (p = 0.161, two-way ANOVA, Fig. 6C). Therefore, there were no significant effects of treatment for skewness or kurtosis in any of the diets. The primary finding was decreased axial vascular complexity with brain maturation as illustrated in Fig. 6J.

**Figure 6.**
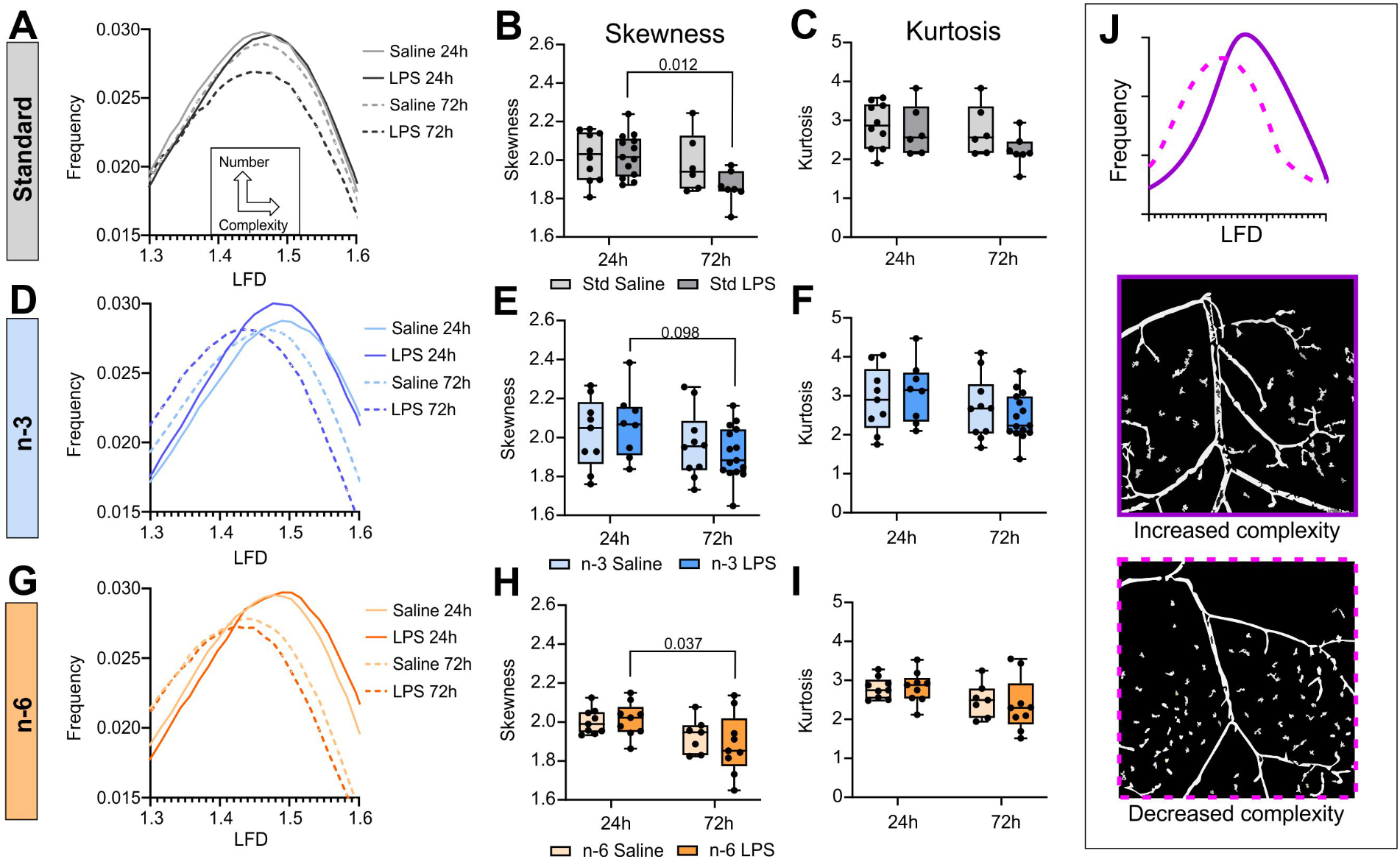
Vessel network complexity is decreased with time but not affected by diet. Vessel network complexity analysis of the axial surface of the brain, based on the distribution of the local fractal dimension of vessels. Results are shown for the standard (A-C), n-3 (D-F), and n-6 (G-I) diets. LFD curves are shown at 24 and 72 h for both saline and LPS conditions (A, D, G). Skewness values of the LFD curves (B, E, H) represent the complexity of the vessel network. Kurtosis values represent the number of vessels (C, F, I). Representation of different complexity levels are shown in (J).

Examining the M3 MCA branch at 72 h after LPS administration revealed two distinct types of staining: 1) well-delineated DiI-positive large arteries as well as capillary beds in some mice, and 2) incomplete/missing capillary bed staining in other mice (Fig. 7A). Mice in each group were dichotomized and Chi^2^ analysis of their proportion did not show significant difference between any of the groups (Fig. 7B). However, mice in the n-3 diet had a 3-fold increased risk to have a decreased capillary bed when treated with LPS (Relative risk = 3.15, Fig. 7B). Whether considering all the samples (with and without capillary bed stain) or only the dichotomized samples, we did not find significant difference between diets or treatments for junction density (Fig. 7C). For total vessel length, there was a significant decrease in the n-3 group treated with LPS compared to the saline group, but this was only true when considering all samples (Sidak’s test: p = 0.029, Fig. 7D). Overall, these results suggest a potential increased sensitivity to LPS in pups from n-3 diet-fed dams.

**Figure 7.**
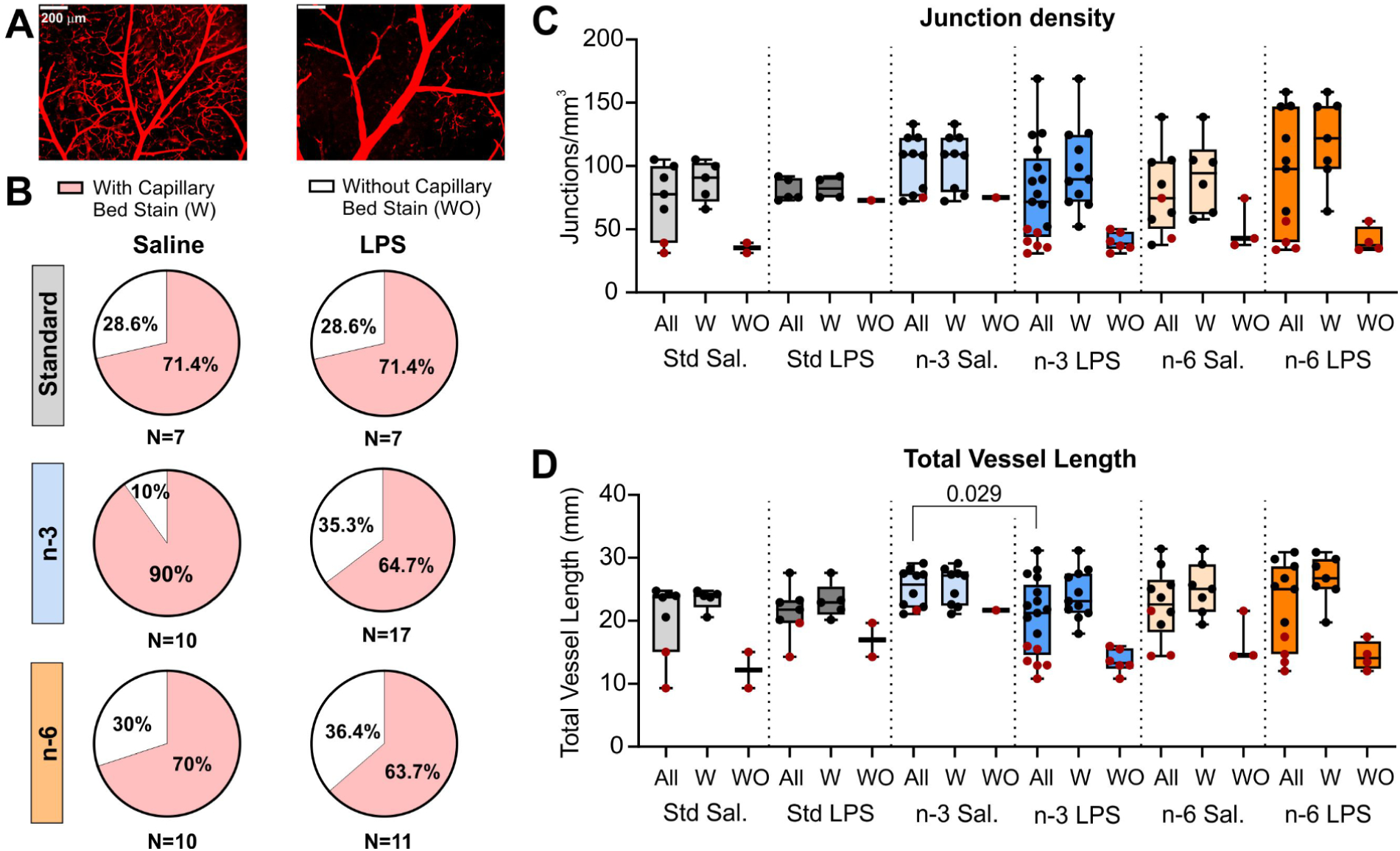
Vessel characteristics within the MCA vascular territory show two populations of mice 72 h after LPS administration. (A) Example of a brain with staining of the capillary bed (left) and without clear staining of the capillary bed (right), scale bar = 200μm. (B) Proportion of mice per group with and without clear staining of the capillary bed at the MCA level is not significantly different between groups but shows more risk for LPS mice in the n-3 diet to have lower capillary stain compared to saline mice (Chi^2^ test and odd ratio). Junction density (C) and total vessel length (D) are shown for all groups, segregating samples with and without capillary bed staining.

## Discussion

We have demonstrated that early-life inflammation following systemic LPS administration in mice at P9 led to a shift in brain microvessel transcriptomics towards a proinflammatory and cell death phenotype at P10 in all diets with higher immune reactivity in PUFA supplemented diets. Concurrently downregulation of cell proliferation and ECM genes triggered by LPS in standard diet group was diminished (n-6) or absent (n-3) in PUFA supplemented diets. While pups from n-3 and n-6 PUFA enriched diets showed the same directionality of LPS-triggered changes in brain vessel transcriptomics, these diets had opposing effects on systemic inflammation where n-3 diet led to faster resolution of inflammation and normalization of cytokine levels and n-6 diet had the opposite effect. At the same time, PUFA diets lowered the levels of cytokines in the brain rather than their response to inflammatory stimulation. However, when analysing the vessel network on the axial surface, neither diet nor LPS treatment affected the vessel morphology and characteristics such as vessel density, junction density and vessel length.

Endotoxin-induced systemic inflammation severely modulated transcriptomics of neonatal brain vasculature, primarily upregulating pathways related to inflammation and innate immune system regulation. The immune role of endothelial cells has gained increasing attention in recent years, including their role in central nervous system inflammation (Amersfoort et al., 2022; Wu, Liu, & Zhou, 2017). Interestingly, we found that supplementation of the diet with PUFA independent on whether they belong to n-3 or n-6 class led to increased immune reaction in brain microvasculature following LPS. Our data showed higher prevalence of upregulation of immune system related genes over downregulation and higher activation scores for cytokine storm, inflammatory pathways and pathways related to blood cell migration, adhesion and binding in both PUFA supplemented diets compared to the standard diet (Fig. 1B, Table 1). Increased reactive state of brain microvessels after LPS injection in animals fed the n-3 diet was not reflected in the cytokine profiles of blood or brain. On the contrary, blood cytokine response to LPS was often the lowest in n-3 diet and the highest in n-6 diet. In brain, lower LPS-stimulated cytokine levels in PUFA supplemented diets could be partially explained by the lower basal levels of cytokines reported previously (Chumak et al., 2022). This discrepancy between the highly upregulated inflammatory signalling in brain microvessels and the low inflammatory status in blood and brain in n-3 diet group may suggest that endothelial cells are not the primary source of cytokines contributing to systemic circulation and brain tissue. It may also depend on the fact that some innate immunity functions are deficient in neonates compared to adults (Kumar & Bhat, 2016).

Our findings often showed same directionality of the effects of PUFA-supplementation independent on prevalence of either n-3 or n-6 fatty acids in the diet. While traditionally n-3 PUFAs were believed to possess anti-inflammatory properties and n-6 PUFAs were considered to be purely pro-inflammatory, this concept has undergone modifications in recent years (Calder, 2006; Djuricic & Calder, 2021; Innes & Calder, 2018). For example, cell culture studies show that both n-3 and n-6 PUFAs can attenuate the baseline release of cytokines or promote/decrease the release of pro-inflammatory cytokines, or supress expression of adhesion molecule and coagulation factors after inflammatory stimulation (Hung et al., 2023; Maucher, Schmidt, Kuhlmann, & Schumann, 2020; Trommer, Leimert, Bucher, & Schumann, 2017). A recent study by Hellstrom et al. reports association of higher DHA blood levels with lower severity of retinopathy of prematurity only when the level of AA in blood is sufficiently high (Hellstrom et al., 2021). These and our findings suggest that the nature of PUFA action on inflammation and neurovasculature during development are more complex than previously thought, at least after infection.

LPS-induced endothelial activation can be accompanied by cell death via apoptosis, necroptosis and pyroptosis (Abdul, Ward, Dong, & Ergul, 2018; Bannerman & Goldblum, 2003; Shioiri et al., 2009; Zhao, Liu, & Chang, 2021). We also observed upregulation of these cell death pathways after LPS in the brain microvessels of the pups, independent of diet. Endothelial cell death has been shown to elevate permeability of the BBB in vitro and in vivo (Banks et al., 2015; Cardoso et al., 2012). However, it has been demonstrated that BBB leakage in vivo occurs mainly at higher doses of LPS than that used in our study (Banks et al., 2015; De Bock et al., 2022; Du & Wang, 2020; You & Jiang, 2021) and the integrity of the BBB in our study was likely unaltered based on microvessel transcriptomics which showed no alteration in regulation of cell junction genes.

Endothelial cell death can prevent angiogenesis (Dimmeler & Zeiher, 2000). Angiogenesis is active in the postnatal brain, expanding the capillary network and reaching stabilization by P15-P25 when proliferation rates of endothelial cells and pericytes decline (Coelho-Santos & Shih, 2020). Consequently, cell death pathways activated by LPS in brain microvessels of animals fed the standard diet, together with altered cell proliferation pathways, may result in brain vascularization deficits that would be prevented in animals fed PUFA-supplemented diets. In addition, changed ECM gene expression observed in LPS-treated standard and n-6 diet groups can impact angiogenesis (Mongiat, Andreuzzi, Tarticchio, & Paulitti, 2016). Furthermore, overrepresentation analysis showed upregulation of angiogenic signalling in the brain microvessels of animals fed n-3 PUFA diet. LPS and inflammation are known to affect angiogenesis (Jeong, Ojha, & Lee, 2021; Pollet et al., 2003). For example, microvascular density in the brain is increased in neurological disorders conditioned by chronic neuroinflammation such as Alzheimer’s disease and multiple sclerosis (Kirk & Karlik, 2003; Vagnucci & Li, 2003). Proinflammatory cytokines such as IL1β, IL6 and TNFα promote endothelial cell proliferation and stimulate angiogenesis (Chen, Chen, Hsiao, Chang, & Chern, 2013). We recently found higher vascularization in the hippocampus of pups 7 days after LPS exposure (Ardalan et al., 2022). A single LPS injection at P4 was also shown to increase retinal vascularization as soon as 48 hours later, eventually resulting in lack of vascularization at P21 (Tremblay et al., 2013). Similarly, prolonged exposure of prematurely born sheep to LPS resulted in decreased vessel density and neurovascular remodelling in the cerebral cortex and white matter (Disdier et al., 2020).

At the same time, there is evidence suggesting that PUFAs can modulate angiogenesis in vitro (Szymczak, Murray, & Petrovic, 2008; Taha, Sharifpanah, Wartenberg, & Sauer, 2020). N-3 PUFA diet has been shown to increase the number of proliferating endothelial cells, length and number of vessels after stroke in aged mice (Cai et al., 2017). In recent neonatal mouse study, we showed that n-3 PUFA supplemented maternal diet decreased brain injury size in a model of neonatal stroke, suggesting a beneficial role in post ischemic inflammation and neuroprotection, while positive effects on neurovascular recovery were not studied and cannot be excluded (Chumak et al., 2022). To our knowledge, no other studies have assessed the effects of PUFA-supplemented maternal diet in the young or adult offspring in terms of cerebral vascular remodelling. However, some groups have shown beneficial effects of PUFA diet in rodents in stroke studies. A 4-week PUFA-enriched diet in adult mice prior to ischemic stroke onset led to better histological, motor and neurological outcomes (Gonzalo-Gobernado et al., 2019). A single dose of DHA injected to adult rats one to four hours after ischemic stroke also showed beneficial effects including decreased lesion volume and behavioural improvement (Ludmila Belayev et al., 2018; L. Belayev et al., 2011). Finally, using transgenic mice overproducing n-3 PUFAs, Wang and colleagues showed a beneficial effect of n-3 PUFAs after transient ischemic stroke with improved revascularization and angiogenesis (Wang et al., 2014).

In terms of vascularization deficits, our vessel painting results did not show any significant effects on vessel or junction density due to LPS treatment. However, at the level of the MCA in the n-3 PUFA diet, we found a decreased total vessel length in LPS animals compared to saline at 72 h. Effects of LPS on signalling pathways might take longer than 72 h to reveal changes in the brain vasculature, which would explain why we did not see any major angioarchitecture differences between diets and treatments.

While we focused on the vascular effects in the offspring of dams fed n-3 enriched or deficient (n-6 supplemented) PUFA diets, deficiency of maternal diet in n-3 PUFAs can affect microglial and oligodendrocyte homeostasis during postnatal brain development even without LPS challenge, as was demonstrated by altered microglial function and enhanced phagocytosis of synapses by microglial cells in P21 mice (Madore et al., 2020) and by disrupted oligodendrocyte maturation and myelin integrity (Leyrolle et al., 2022).

Our study has some limitations: as mentioned above, we only examined effects of diets and LPS up to 72 h after treatment. The study would also benefit from the cytokine level analysis at the shorter time point to describe the early stages of inflammation. While we focused on mouse pups, we believe it would be interesting to study effects at later time points in young adults and adults. It is known that maternal diet during foetal development has life-long lasting effects on offspring (Godfrey & Barker, 2001; Langley & Jackson, 1994). It should also be noted that the cytokine and gene assays were performed on whole cerebrum purified microvessels while the vessel painting experiments only assessed the vascular network on the axial cortical surface. Hence, we are not comparing the exact same population of microvessels. However, we expect the systemic effects of LPS to affect all brain vessels the same way.

## Conclusion

The current study shows that PUFAs content in maternal diet can modulate not only inflammatory response but also functional response of offspring brain vasculature to neonatal inflammation and thus contribute to long-term neurodevelopmental consequences. At that, our results challenge commonly accepted notion of opposing biological effects of PUFAs from n-3 and n-6 classes, at least in the neonatal infection-related model, and highlight the importance of maternal diet during pregnancy and lactation.

## Acknowledgements

We thank Novogene Co., LTD (Cambridge, UK) for performing the sequencing and Katarine Tuvé (Genomics and Bioinformatics Core Facility platforms at the Sahlgrenska Academy, University of Gothenburg) for performing the read alignment and expression quantification of the sequencing raw data. We acknowledge Kajsa Groustra (Experimental Biomedicine, Sahlgrenska Academy, University of Gothenburg) for help with mouse breeding and feeding the diets and Anna-Lena Leverin (Institute of Neuroscience and Physiology, Centre of Perinatal Medicine and Health, Sahlgrenska Academy, University of Gothenburg) for technical support. We thank Mary Hamer and Kara Wendel (Department of Pediatrics, University of California Irvine, Irvine CA, USA) for help with mouse breeding and perfusion for the vessel painting experiments. Graphical abstract was created using BioRender.com.

## Author contribution

TC, AJ, CM, JE, AO and ZW conceptualized the study, designed experiments and wrote the manuscript. TC, AJ, MA, JE, PS, RQ, AS and SS performed animal studies, laboratory experiments, data processing and analysis. All authors contributed to data interpretation and critical revision of the manuscript.

## Supplemental figures

**Suppl. Figure 1. N-6 diet affects post-LPS weight gain but not brain area in the offspring.** Effect of systemic LPS administration on the weight gain in pups fed different diets. (A) Weight gain 24h after LPS or saline injection in pups from dams fed different diets, t-test. (B) Weight gain 72h after LPS or saline injection in pups from dams fed different diets, t-test. (C) Weight of pups from dams fed different diets at P9, at the time of LPS or saline injection. Welch’s ANOVA, Dunnett’s post-hoc test. (D) Axial brain area 24 h after injection. (E) Axial brain area 72 h after injection.

**Suppl. Figure 2. Gene expression analysis suggests intact blood-brain barrier in all diets and a stronger pro-angiogenic response to LPS in n-3-supplemented diet group.** PCA plots of individual genes annotated to cell junction and angiogenesis regulated by LPS within each diet. Variables are colour coded according to distance between them rendering upregulated genes red and downregulated genes blue. Numbers of regulated genes are stated in upper right corner out of all analysed genes for each process (in brackets).

**Suppl. Figure 3. PUFA diets modulate cytokine levels 72 h after LPS injection.** Cytokine levels in the offspring blood plasma and brain 72 h after LPS injection. (A) Heatmap of cytokine levels analysed in blood plasma and brain of offspring from dams fed the standard, the n-3 or the n-6 enriched diet and injected with saline or LPS at P9. Each column refers to one animal and each row shows levels of a specific cytokine. Each cytokine level was normalized over all samples using Z-score normalization and is shown using a colour scheme based on Z-score distribution from -2 to 2. Cytokines regulated by LPS (based on two-way ANOVA results) are highlighted with the frame. (B, C) Graphs showing the reaction to LPS in plasma (B) or brain (C). Zero level indicates no difference between the level of the cytokine in LPS sample and the mean level of the cytokine among the saline samples. Only graphs with significant differences between diet groups are presented. P values presented on the graphs show results of the post-hoc test. One-way ANOVA with Fisher’s LSD (B:IL4, CCL19, CCL24, CCL7, CCL20; C: CCL22) or Kruskal-Wallis with uncorrected Dunn’s (B: CCL22, TNFα, CCL17; C: CCL11) post hoc tests.

## Supplemental Tables

**Supplemental Table 1.** Z-scores for selected top LPS regulated canonical pathways and biological functions in standard diet with comparisons to other diets generated by IPA.

**Supplemental Table 2.** Pathways enriched by LPS across diets generated by REACTOME.

**Supplemental Table 3.** Descriptive statistics and results of two-way ANOVA analysis for cytokine levels in blood plasma of pups from different diet groups 24 h after LPS injection.

**Supplemental Table 4.** Descriptive statistics and results of two-way ANOVA analysis for cytokine levels in blood plasma of pups from different diet groups 72 h after LPS injection.

**Supplemental Table 5.** Descriptive statistics and results of two-way ANOVA analysis for cytokine levels in brain of pups from different diet groups 24 h after LPS injection.

**Supplemental Table 6.** Descriptive statistics and results of two-way ANOVA analysis for cytokine levels in brain of pups from different diet groups 72 h after LPS injection.

